# Functional Activity, Functional Connectivity and Complex Network Biomarkers of Progressive Hyposmia Parkinson’s Disease with No Cognitive Impairment: Evidences from Resting-state fMRI Study

**DOI:** 10.1101/2024.05.22.595435

**Authors:** Lei Geng, Wenfei Cao, Juan Zuo, Hongjie Yan, Jinxin Wan, Yi Sun, Nizhuan Wang

## Abstract

**Background:** Olfactory dysfunction stands as one of the most prevalent non-motor symptoms in the initial stage of Parkinson’s disease (PD). Nevertheless, the intricate mechanisms underlying olfactory deficits in Parkinson’s disease still remain elusive.

**Methods:** This study collected rs-fMRI data from 30 PD patients (15 with severe hyposmia (PD-SH) and 15 with no/mild hyposmia (PD-N/MH)) and 15 healthy controls (HC). To investigate functional segregation, the amplitude of low-frequency fluctuation (ALFF) and regional homogeneity (ReHo) were utilized. Functional connectivity (FC) analysis was performed to explore the functional integration across diverse brain regions. Additionally, the graph theory-based network analysis was employed to assess functional networks in PD patients. Furthermore, Pearson correlation analysis was conducted to delve deeper into the relationship between the severity of olfactory dysfunction and various functional metrics.

**Results:** Firstly, PD patients showed significantly higher ALFF values in the superior temporal gyrus compared to HC, especially in the PD-SH group versus PD-N/MH. Meanwhile, ALFF values in this region negatively correlated with olfactory testing scores. Secondly, PD patients had higher ReHo in middle temporal gyrus and superior frontal gyrus than HC, especially PD-SH. Meanwhile, olfactory scores negatively correlated with these ReHo values. Thirdly, we observed a negative correlation between superior cerebellar-insula connectivity and olfactory scores, suggesting a neural circuit link to olfactory dysfunction. Lastly, the PD patients’ brain networks consistently showed small-world attributes, with the PD-N/MH group having significantly higher nodal betweenness in the superior cerebellum than the PD-SH group. There’s also a positive link between superior cerebellum’s betweenness and olfactory scores. Additionally, there is a notable difference in the node degree of the superior temporal gyrus when comparing the PD-SH group to the PD-N/MH group.

**Conclusion:** Using fMRI, our study analyzed brain function in PD-SH, PD-N/MH, and HC groups, revealing impaired segregation and integration in PD-SH and PD-N/MH. We hypothesize that changes in temporal, frontal, occipital, and cerebellar activities, along with aberrant cerebellum-insula connectivity and node degree and betweenness disparities, may be linked to olfactory dysfunction in PD patients.

## 1. Introduction

Parkinson’s Disease (PD), whose impact on society is significant, is a common neurodegenerative disease characterized by motor dysfunction, tremor, muscle rigidity, and non-motor symptoms [1,2]. With population growth and aging, the epidemiology of Parkinson’s disease has changed considerably, and its incidence rate is increasing. A recent Global Burden of Disease (GBD) study showed that age-standardized incidence rates of Parkinson’s disease increased largely among nervous system diseases [3]. However, the causes of Parkinson’s disease are complex and uncertain, and current researches suggest that genetics, environment, and interactions thereof may be involved [4,5,6,7,8]. In clinical settings, Parkinson’s disease is primarily diagnosed through a comprehensive assessment of medical history and neurological symptoms [9]. It’s worth noting that when clinical manifestations appear, at least 50%-60% of dopaminergic neurons have already died, which can potentially delay treatment [10]. The topic of motor symptoms, including tremor, rigidity, and slowness, is often crucial and cannot be overlooked. In addition to the motor features, there are also many non-motor features, such as olfactory dysfunction, cognitive decline, sleep disorders, depression and apathy [11,12]. Olfactory abnormalities usually appear before motor symptoms, known as “prodromal” symptoms, which means that changes in the olfactory pathway may be the earliest pathological changes in PD [13].

As a subjective sensation, olfaction is inherently a chemical sensing process that involves the detection of specific odor molecules by sensory cells [14]. The olfactory system comprises the peripheral olfactory system, such as the olfactory neuroepithelium and olfactory tract, as well as the central olfactory system, including the olfactory bulb and olfactory cortex [15]. The specific binding of odor ligands to receptors on olfactory neurons triggers the activation of the second messenger pathway, resulting in the generation of action potentials. Subsequently, these action potentials are transmitted in a graded fashion through the olfactory tract, ultimately reaching the olfactory bulb and olfactory cortex. This sequential process constitutes the fundamental mechanism underlying the production of olfactory sensation [16, 17]. A fully functional olfactory system enables humans to assess the safety of consumed food and perceive vital information about surrounding dangers. However, this capability gradually diminishes as individuals age [18]. Olfactory dysfunction, characterized by the diminution or absence of olfactory function, can be congenital, idiopathic, or associated with cognitive disorders [19–22]. Among the prevalent causes of this condition are sinus diseases, infections, and head trauma [23]. Olfactory impairment stands as the most prevalent non-motor symptom among individuals with PD, affecting over 90% of patients who exhibit notable olfactory decrement [24]. The impairment of odor recognition has been recognized as a pivotal central defect that characterizes both PD and Alzheimer’s disease (AD) [25]. In addition, olfactory function testing can assist in the early diagnosis of PD, so a large number of researchers have attempted to focus on the study of olfactory dysfunction in PD patients [26, 27]. However, the intricacies surrounding the mechanism of olfactory impairment in PD patients remain elusive, necessitating deeper investigation through rigorous molecular, pathological, and imaging research endeavors.

Magnetic Resonance Imaging (MRI) has been extensively employed to investigate the structural, functional, and metabolic changes in the brain linked to olfactory dysfunction in PD patients, thus facilitating the search for diagnostic biomarkers [28–33]. Resting-state functional magnetic resonance imaging (rs-fMRI) is a widely utilized imaging technique for evaluating the dynamic alterations in brain function among patients in a resting state. Amplitude of low-frequency fluctuation (ALFF) [34], Regional Homogeneity (ReHo) [35], functional connectivity (FC) [36], and complex network analysis serves as integral components of the evaluation indicators [37–38], enabling the exploration of alterations in brain function from both the angles of functional segregation and functional integration.

Therefore, leveraging functional magnetic resonance imaging (fMRI) technology, this study aims to comprehensively investigate the brain functional alterations in PD patients exhibiting severe olfactory dysfunction yet maintaining normal cognitive function. By employing a range of indicators from diverse perspectives, we seek to further analyze the correlation between abnormal neuroimaging markers and olfactory scores, ultimately unraveling the potential neuroimaging mechanisms underlying olfactory impairment in PD patients.

## 2. Materials and Methods

### 2.1 Participants and Dada acquisition

In this research, we utilized a publicly accessible dataset sourced from the OpenfMRI database (https://openfmri.org/dataset/ds000245/). The accession number is ds000245, and the original article contains comprehensive information regarding both the demography and clinical data [39]. Patients who were diagnosed with PD at the Department of Neurology at Nagoya University were included in this study. All participants underwent the Japanese Odor Stick Identification Test (OSIT-J) [40] and the Addenbrooke’s Cognitive Examination-Revised (ACE-R) [41] to evaluate their odor-identification ability and cognitive function, respectively. In conclusion, 15 PD patients with severe hyposmia (PD-SH) but no cognitive impairment and 15 PD patients with no/mild hyposmia (PD-N/MH) and no cognitive impairment were included in this study. Additionally, 15 healthy subjects without cognitive impairment, hyposmia, and family history of PD were enrolled as healthy controls (HCs). For more detailed information, kindly refer to the reference [39]. All subjects provided their explicit informed consent by signing the necessary agreement, indicating their willingness to participate in the study. This study received the formal approval of the Ethical Committee of Nagoya University Graduate School of Medicine.

All MRI and functional magnetic resonance imaging (fMRI) scans were executed employing a 3.0T scanner (Siemens, Erlangen, Germany) with a 32-channel head coil at Nagoya University’s Brain and Mind. The high-resolution T1-weighed images were obtained with following standards: repetition time (TR) = 2.5 s, echo time (TE) = 2.48 ms, 192 sagittal slices with 1-mm thickness, field of view (FOV) = 256 mm, matrix=256 × 256. The resting-state fMRI parameters in this study were carefully controlled to adhere to the following standards: TR = 2.5 s, TE = 30 ms, 39 transverse slices with a 0.5-mm inter-slice interval and 3-mm thickness, FOV = 192 mm, matrix=64 × 64, flip angle = 80 degrees.

### 2.2. FMRI Data Preprocessing

Before commencing data analysis, the rs-fMRI data underwent a series of meticulous preprocessing procedures and rigorous quality control measures. The tasks were carried out meticulously using the Data Processing and Analysis of Brain Imaging (DPABI) toolbox (version 6.1, http://rfmri.org/dpabi), which is based on MATLAB (The Math Works, Natick, MA, USA) software. This toolbox was utilized to eliminate the effects of data acquisition, physiological noise, and individual variations in the subject’s brain [42]. Additionally, it ensured the reliability and sensitivity of the group-level analysis. The preprocessing steps are detailed as follows: (1): The fMRI data originally in DICOM format underwent conversion to 3D-NIFTI format, leveraging the operating environment provided by MATLAB R 2021a. (2): To minimize errors caused by unstable initial environments during image acquisition, we have removed the first 10 time points. (3): The time difference inherent in fMRI images, resulting from compartmentalized scanning, which causes images to be acquired at different times, was rectified. (4): To correct small head movements, head motion correction was utilized. The exclusion thresholds were set at 2 mm and 2°, meaning that any movement exceeding 2 mm in any direction or a rotation exceeding 2° in any direction would be excluded. (5): To ensure comparability among varying individual images, the raw images of each subject were precisely aligned to the standard MNI space (featuring a resolution of 3 mm × 3 mm × 3 mm). (6): To enhance the signal-to-noise ratio, minimize registration errors, eliminate poorly registered image data, we employ Gaussian smoothing with a 4mm Full Width Half Maximum (FWHM). (7): The low-frequency band of the BOLD signal offers a more accurate representation of spontaneous neural activities in the human brain during physiological states. Therefore, in this study, we employed high-pass filtering to select the BOLD signal within the frequency range of 0.01-0.1 Hz.

### 2.3. ALFF Analysis

Before high-pass filtering was applied, the ALFF was computed using the DPABI software [43], utilizing the preprocessed dataset. This involved converting the time series data of each individual voxel into a frequency spectrum using the Fourier transform. Then the amplitudes within the frequency range of 0.01-0.1 Hz were summed to calculate the ALFF [44–46]. The analysis of variance (ANOVA) was conducted for ALFF in three groups, and the Turkey honest significant difference (HSD) method was used to correct the errors between pairwise comparisons, with age and gender as covariates. A rigorous correction for multiple comparisons was applied at the clump level in this study to minimize the false-positive rates. And the present study then reveals the surviving corrected clumps.

### 2.4. ReHo Analysis

We used the DPABI software to perform ReHo analysis based on preprocessed data (prior to smoothing). The ReHo map of the subject was obtained by calculating the Kendall’s coefficient of concordance between the time series of each voxel and the time series of the surrounding 26 voxels in a voxel-wise manner [47]. Next, the obtained ReHo map still needs to undergo spatial smoothing to improve the signal-to-noise ratio and correct registration errors caused by normalization. The statistical approach for calculating ReHo is identical to the statistical analysis method utilized for calculating ALFF as described earlier. Similarly, we extracted the ReHo values from PD patients.

### 2.5. Complex Network Analysis

The DPABI NET (version 1.1) toolbox was utilized for brain network construction and subsequent analysis [48]. The entire brain was partitioned into a total of 116 distinct brain regions using the widely recognized brain atlas, the Anatomical Automatic Labeling (AAL) template (http://www.gin.cnrs.fr/en/tools/aal/). The Brain Connectivity Toolbox (https://www.nitrc.org/projects/bct/) [49] was utilized to conduct complex-network analysis, while the BrainNet viewer (https://www.nitrc.org/projects/bnv/) was employed to visualize the results [50]. The analysis of brain network topology encompasses both “small world” properties and node properties. In our study, we calculated the magnitudes of various small-world property across 50 distinct sparsity thresholds. Furthermore, we ascertained the area beneath the curve (AUC) for each small-world parameter within the 0.01-0.34 sparsity interval [51], facilitating a more precise identification of changes in the brain’s small-world architecture. Afterwards, we calculated the values for the following small-world metrics and node properties at each sparsity level: characteristic shortest path length (L_p_), clustering coefficient (C_p_), normalized clustering coefficient (γ), normalized characteristic shortest path length (λ), small-worldness (σ), local efficiency (E_loc_), global efficiency (Eg_lob_), degree centrality, betweenness centrality [52].

### 2.6 Correlation Analysis

Using the DPABI software, we extracted the ALFF, ReHo and FC values for each PD patients (PD patients with severe hyposmia and PD patients with no/mild hyposmia) and subsequently performed a correlation analysis with their OSIT-J scores. During this analysis, we accounted for potential confounding effects by regressing out age and gender as covariates. The statistical significance threshold was established at P < 0.005. Furthermore, we performed a correlation analysis to assess the relationship between the brain network indicators of PD patients and OSIT-J scores, with a significance level of P<0.05 being deemed statistically meaningful.

## 3. Results

### 3.1. Results of ALFF Analysis

#### 3.1.1. PD-SH V.S. HC

The PD-SH group showed that the ALFF value of right inferior cerebellum (AAL: Cerebelum_Crus2_R), left inferior temporal gyrus, left middle temporal gyrus and right temporal pole: superior temporal gyrus was significantly higher (FDR correction method with a significance threshold of P<0.05) than those of the HC group (Figure 1). The ALFF of PD-SH group in bilateral putamen, right rolandic operculum, right insula, bilateral precuneus, right posterior cingulate gyrus, right superior frontal gyrus, orbital part, right gyrus rectus, right olfactory cortex, right hippocampus, bilateral cerebellum (AAL: Cerebelum_8), right inferior occipital gyrus, right inferior frontal gyrus, orbital par, left middle frontal gyrus, orbital part, right lingual, right thalamus, left posterior cingulate gyrus and right rolandic operculum is lower than HC group (Figure 1).

**Figure 1.**
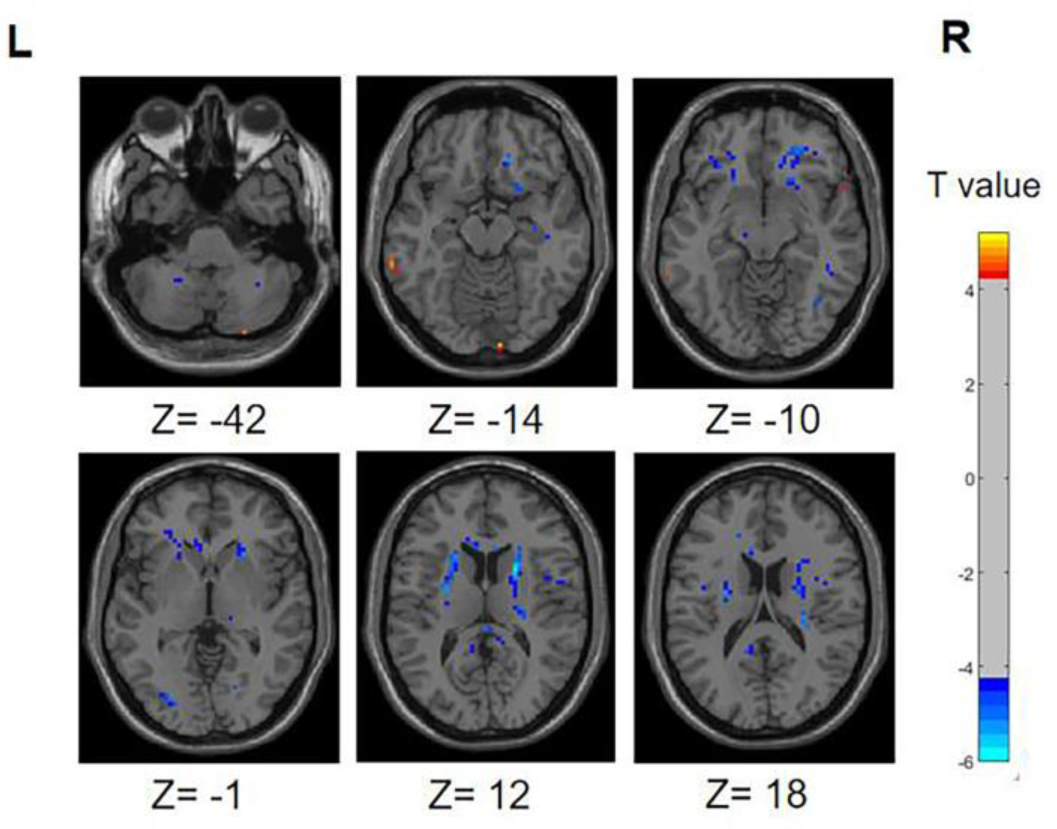
Differences in ALFF between PD-SH group and HC group. The color red denotes increased ALFF in PD-SH group compared to HC group, while color blue denotes decreased ALFF in PD-SH group compared to HC group. The corresponding color bar denotes the t-value. R: right; L: left.

#### 3.1.2. PD-N/MH V.S. HC

Compared with the HCs, the ALFF in the bilateral temporal pole: superior temporal gyrus of the the PD-N/MH group significantly increased (FDR correction, P<0.01). As shown in Figure 2, the ALFF of PD-N/MH group in the left middle occipital gyrus, right putamen, right caudate, left posterior cingulate gyrus and right median cingulate and paracingulate gyri are lower than HC group (FDR correction, P<0.01).

**Figure 2.**
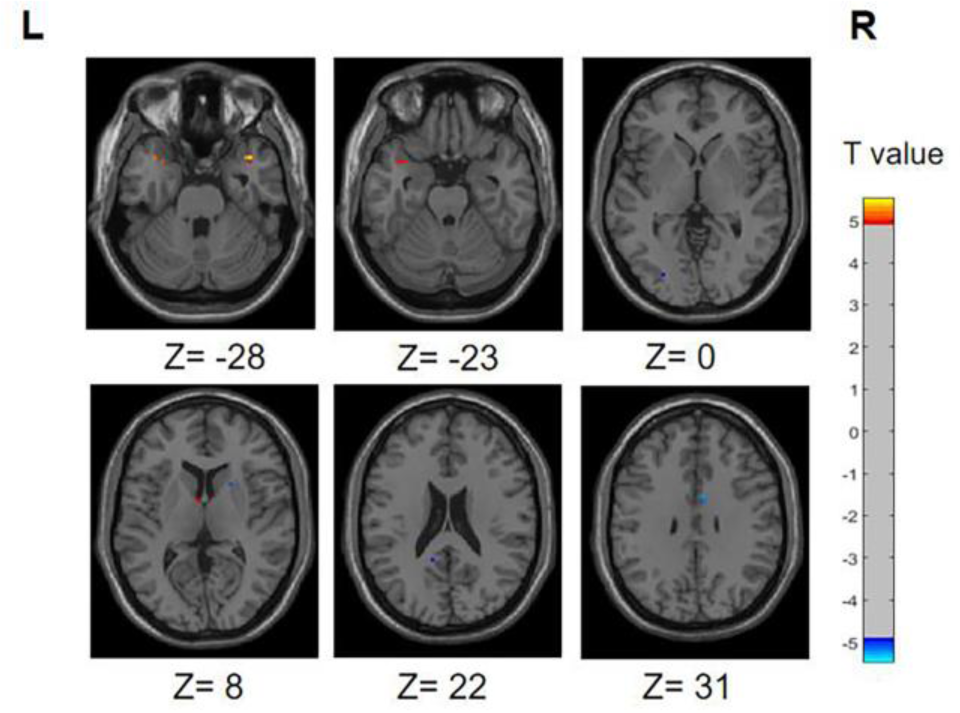
Differences in ALFF between PD-N/MH group and HC group. The color red denotes increased ALFF in PD-N/MH group compared to HC group, while color blue denotes decreased ALFF in PD-N/MH group compared to HC group. The corresponding color bar denotes the t-value. R: right; L: left.

#### 3.1.3. PD-N/MH V.S. PD-SH

Illustrated in Figure 3, the PD-SH exhibited a noteworthy enhancement in ALFF within bilateral superior temporal gyrus, bilateral middle temporal gyrus, left superior cerebellum, left superior frontal gyrus, dorsolateral and inferior frontal gyrus, triangular part, when compared to the PD-N/MH group (voxel p < 0.001, cluster p-value < 0.05, GFR corrected). The ALFF of PD-SH group in left inferior cerebellum, vermis and left anterior cingulate and paracingulate gyri are lower than PD-N/MH group (voxel p < 0.001, cluster p-value < 0.05, GFR corrected).

**Figure 3.**
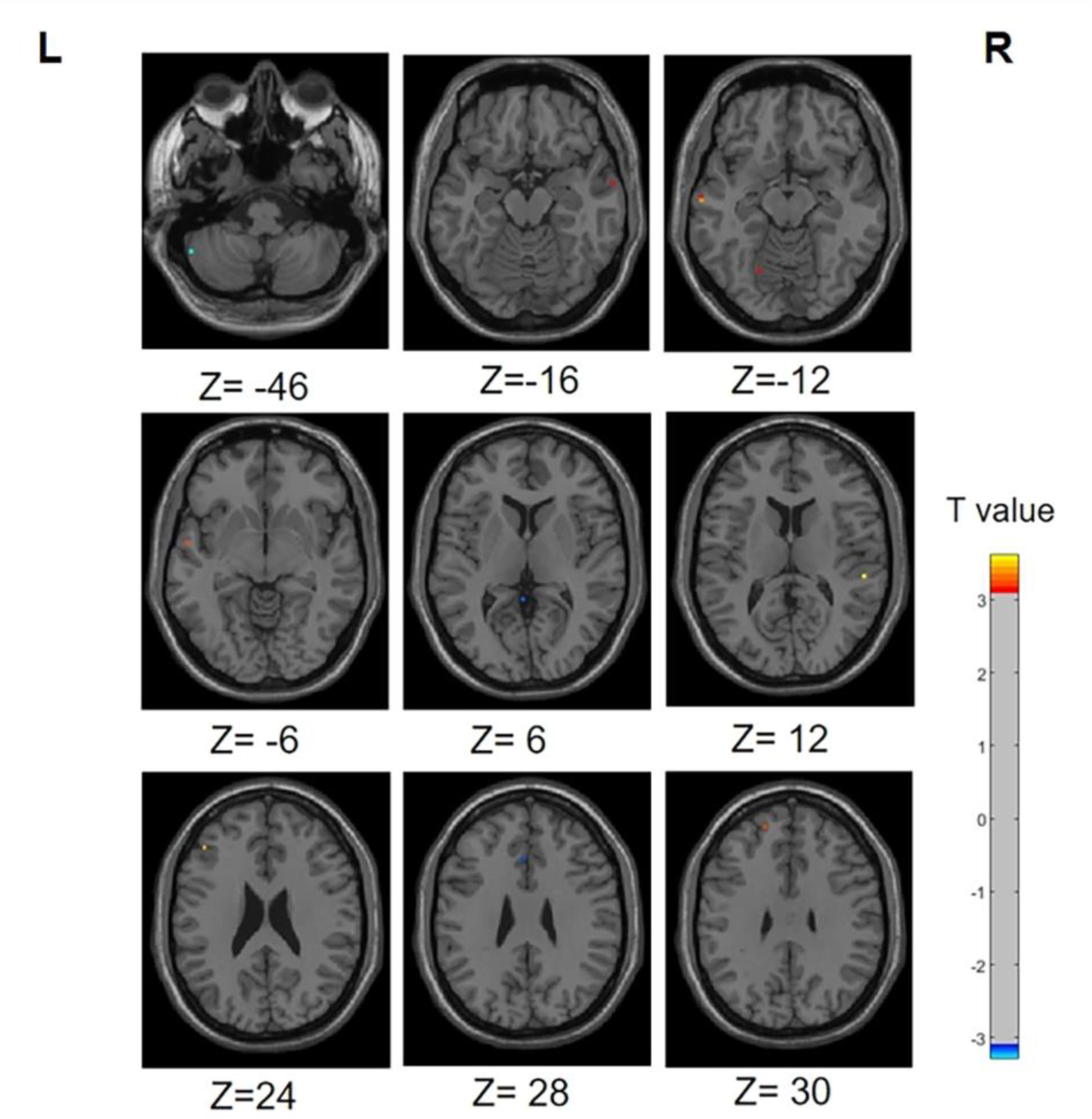
Differences in ALFF between PD-N/MH group and PD-SH group. The color red denotes increased ALFF in PD-SH group compared to PD-N/MH group, while color blue denotes decreased ALFF in PD-SH group compared to PD-N/MH group. The corresponding color bar denotes the t-value. R: right; L: left.

### 3.2. Results of ReHo Analysis

#### 3.2.1. PD-SH V.S. HC

In the PD-SH group, a statistically significant elevation (FDR correction method with a significance threshold of P<0.05) of ReHo was observed in region of left Inferior Cerebellum, left Middle temporal gyrus, right Lingual gyrus, right Inferior frontal gyrus, triangular part, right Middle frontal gyrus, and bilateral Superior frontal gyrus, dorsolateral (Figure 4), when compared to the HC group. In comparison to the HC group, the PD-SH group demonstrated decreased (FDR correction, P<0.05) ReHo in specific regions, including (AAL: Cerebelum_9), bilateral Fusiform gyrus, bilateral Hippocampus, left Middle temporal gyrus, left Inferior temporal gyrus, right Inferior occipital gyrus, left Superior occipital gyrus, right Thalamus, left Precuneus, left Cuneus, right Caudate nucleus and left Lenticular nucleus, putamen.

**Figure 4.**
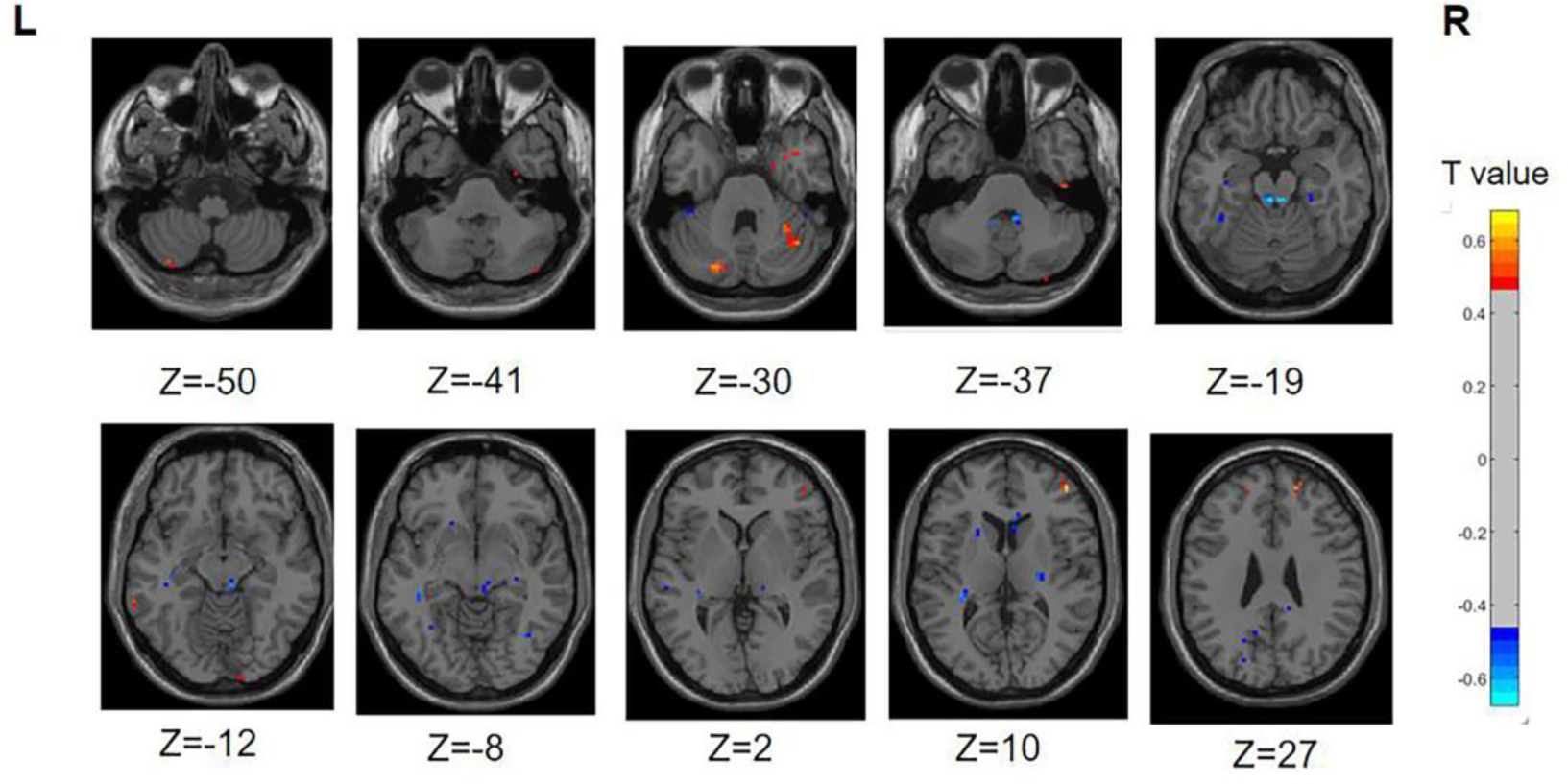
Differences in ReHo between PD-SH group and HC group. The color red denotes increased ReHo in PD-SH group compared to HC group, while color blue denotes decreased ReHo in PD-SH group compared to HC group. The corresponding color bar denotes the t-value. R: right; L: left.

#### 3.2.2. PD-N/MH V.S. HC

The ReHo values in region of bilateral Inferior Cerebellum (AAL: Cerebelum_Crus2), right Superior frontal gyrus, medial orbital, and right Superior frontal gyrus, dorsolateral were significantly higher (FDR correction, P<0.05) in the PD-N/MH group compared to the HC group. Additionally, in the PD-NMH group, lower (FDR correction, P<0.05) ReHo values were observed in regions right Fusiform gyrus, left Superior temporal gyrus, left Lenticular nucleus, putamen, bilateral Posterior cingulate gyrus, right Caudate nucleus and left Precental gyrus compared to the HC group (Figure 5).

**Figure 5.**
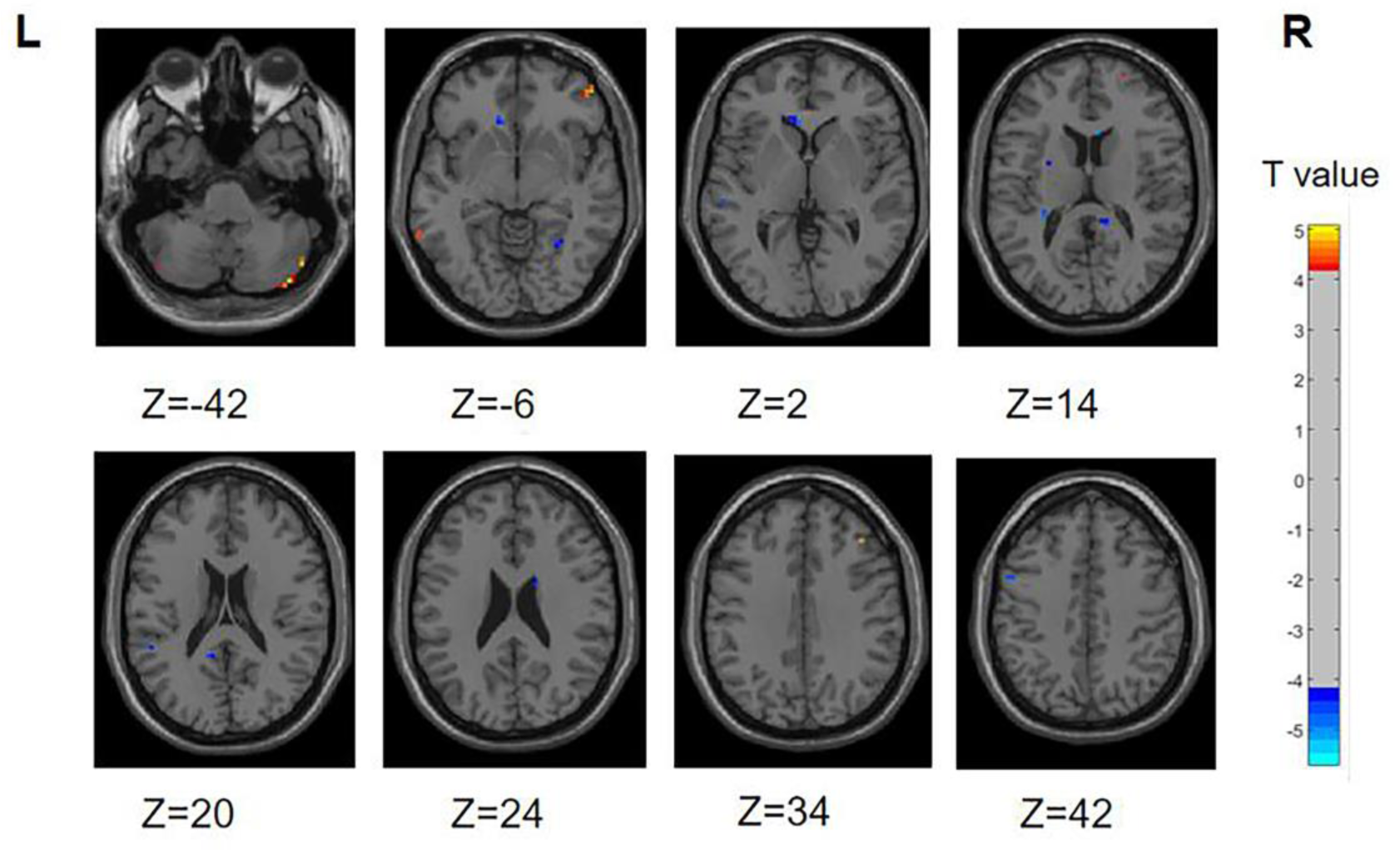
Differences in ReHo between PD-N/MH group and HC group. The color red denotes increased ReHo in PD-N/MH group compared to HC group, while color blue denotes decreased ReHo in PD-N/MH group compared to HC group. The corresponding color bar denotes the t-value. R: right; L: left.

#### 3.2.3. PD-N/MH V.S. PD-SH

Compared to the PD-SH group, the PD-N/MH group exhibited decreased (voxel level p < 0.001, cluster p-value < 0.05, GFR corrected) ReHo in the following regions, including the left Precuneus, left fusiform, left Superior Cerebellum, left Superior frontal gyrus, medial, right Superior frontal gyrus, and right Middle occipital gyrus. Furthermore, our findings revealed that PD-N/MH patients displayed a notable enhancement (voxel p < 0.001, cluster p-value < 0.05, GFR corrected) in ReHo within the right Middle frontal gyrus, left Hippocampus and right Superior frontal gyrus, orbital part in comparison to PD-SH patients (Figure 6).

**Figure 6.**
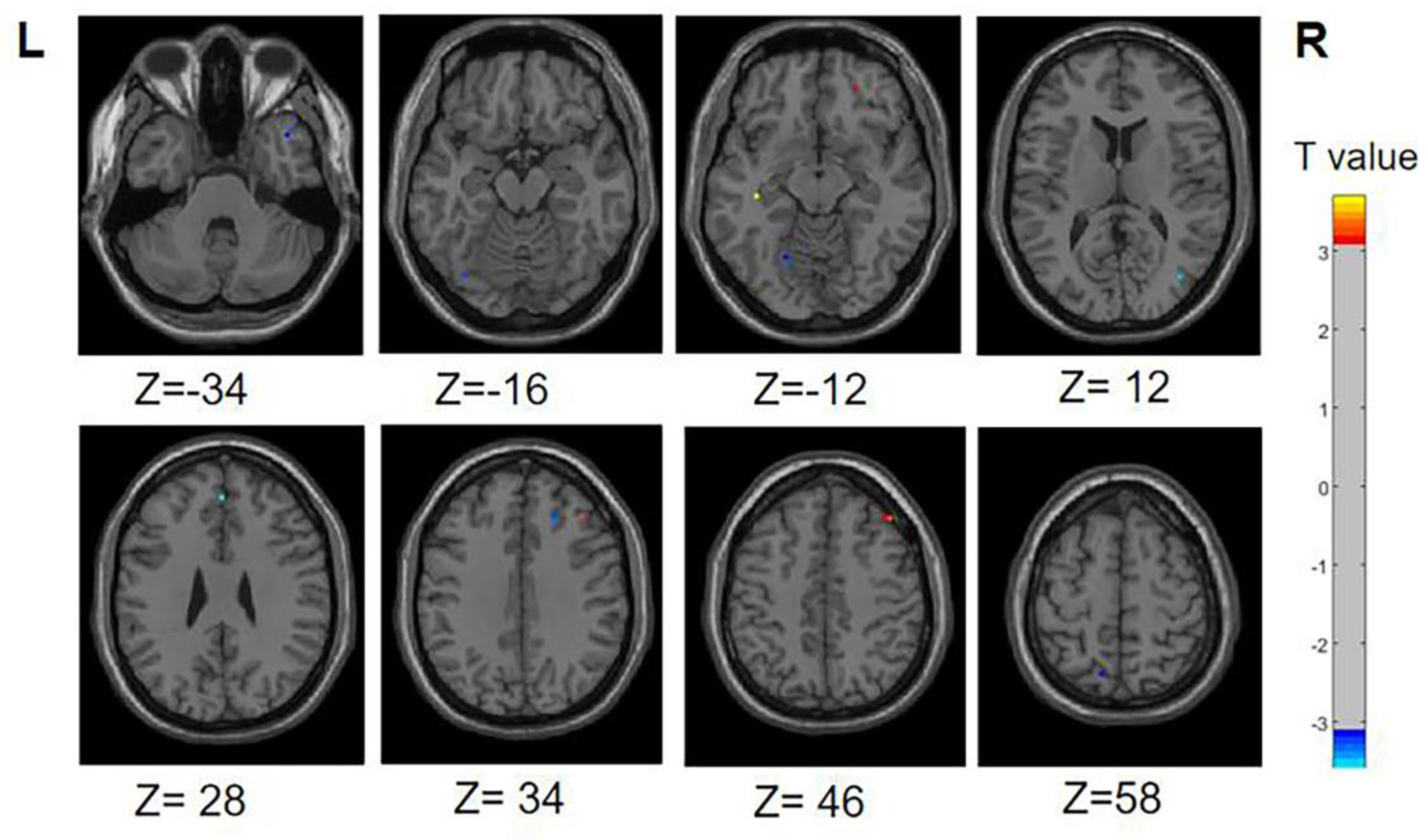
Differences in ReHo between PD-N/MH group and PD-SH group. The color red denotes increased ReHo in PD-N/MH group compared to PD-SH group, while color blue denotes decreased ReHo in PD-N/MH group compared to PD-SH group. The corresponding color bar denotes the t-value. R: right; L: left.

### 3.3. Results of Functional Connectivity Analysis

#### 3.3.1. PD-SH V.S. HC

Illustrated in Figure 7, We observed that the PD-SH group exhibited decreased connectivity in specific regions (FDR correction, P<0.00001), including the connection between the left middle frontal gyrus and left precental gyrus, as well as the link between the left insula and left caudate nucleus, when compared to the HC group. Furthermore, there are regions where connectivity is notably strengthened, including the connection between the left superior frontal gyrus, dorsolateral and left superior frontal gyrus, orbital part, the connection between the left superior frontal gyrus, medial and the left superior frontal gyrus, orbital part, as well as the connection between the left inferior cerebellum and the left superior frontal gyrus, orbital part (FDR correction, P<0.00001).

**Figure 7.**
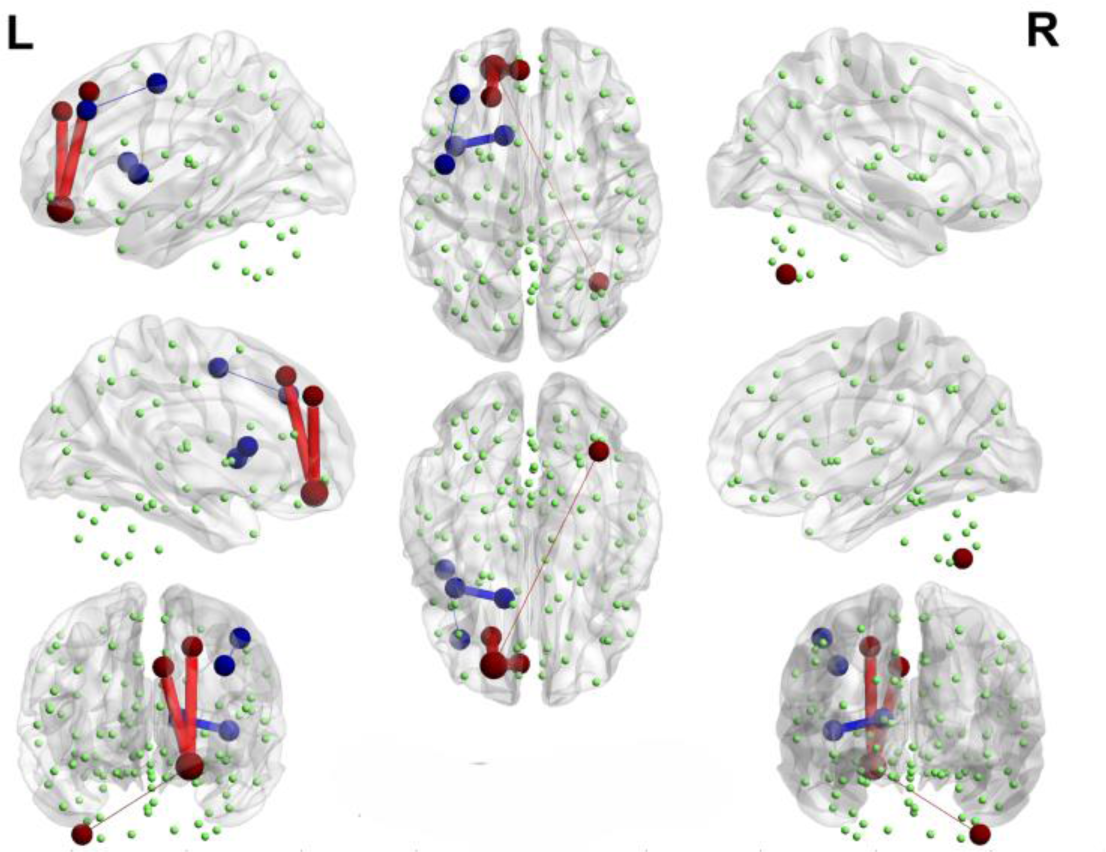
Functional connectivity alterations between the PD-SH group and the HC group. The spheres represent brain nodes, where red spheres and connecting lines signify strengthened functional connectivity, while blue indicates a decrease in functional connectivity. R: right; L: left.

#### 3.3.2. PD-N/MH V.S. HC

Interestingly, when comparing the PD-N/MH group to the HC group, we also observed similarly decreased functional connectivity regions (FDR correction, P<0.00001), particularly in the connections between the left middle frontal gyrus and the left precental gyrus, and connections between the left insula and left caudate nucleus (Figure 8). In addition, when compared to the HC group, the PD-N/MH group demonstrated regions with notably increased connectivity, specifically in the connections between the left superior frontal gyrus, dorsolateral and left superior frontal gyrus, orbital par, the left superior frontal gyrus, medial and the left superior frontal gyrus, orbital part, the left inferior cerebellum and the left superior frontal gyrus, orbital part, the left superior frontal gyrus and the left middle frontal gyrus as well as the left precental gyrus and the right inferior cerebellum (FDR correction, P<0.00001).

**Figure 8.**
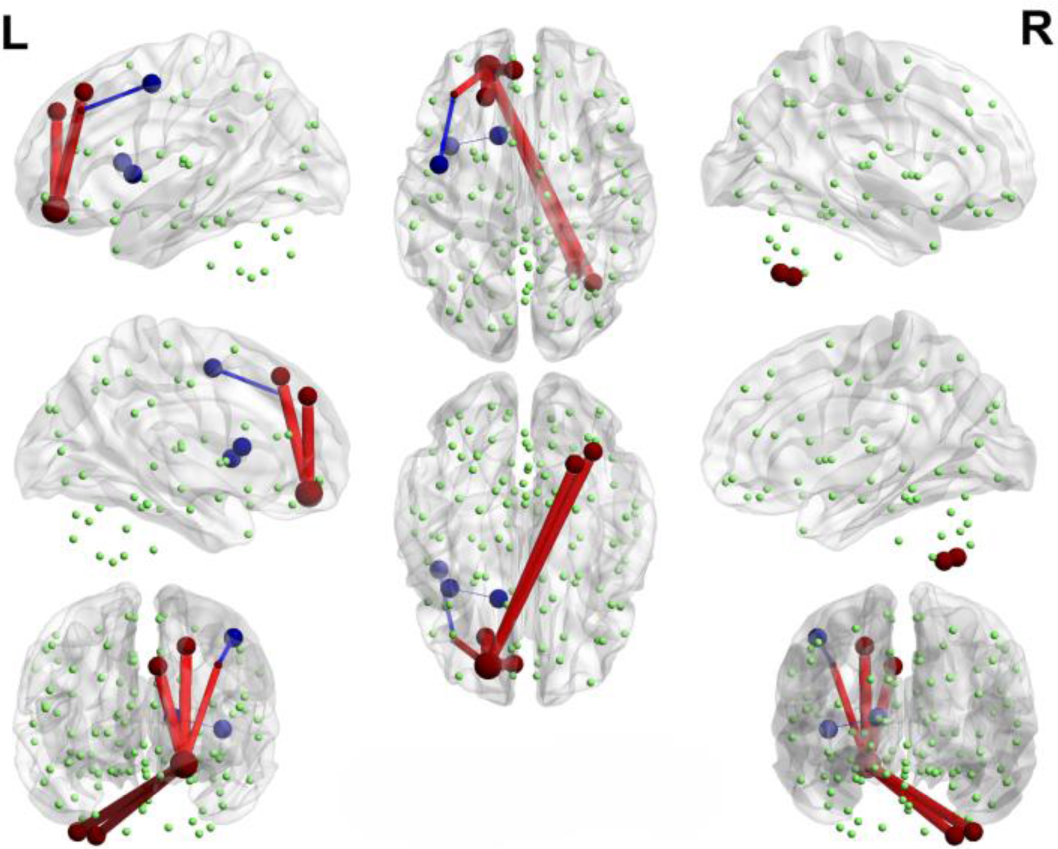
Functional connectivity alterations between the PD-N/MH group and the HC group. The spheres represent brain nodes, where red spheres and connecting lines signify strengthened functional connectivity, while blue indicates a decrease in functional connectivity. R: right; L: left.

#### 3.3.3. PD-SH V.S. PD-N/MH

We conducted a comparison of brain functional connectivity strength between the PD-SH group and the PD-N/MH group (Figure 9), revealing that the PD-SH group demonstrated significantly decreased connectivity between the right superior cerebellum and the vermis (P<0.05).

**Figure 9.**
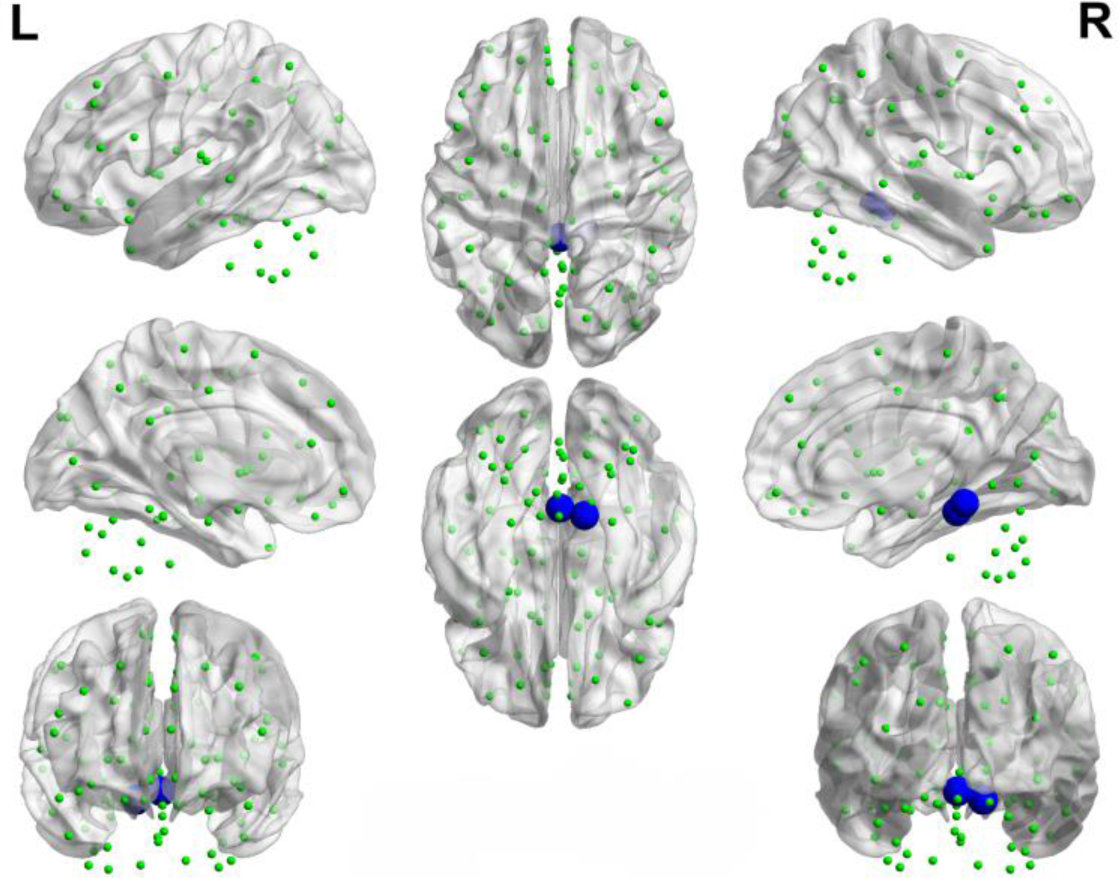
Decreased functional connectivity between the right superior cerebellum and the vermis of the PD-SH group in comparison to the PD-N/MH group. The blue spheres and connecting lines signify decreased functional connectivity. R: right; L: left.

### 3.4. Results of Complex Network Analysis

#### 3.4.1 Small-worldness

Our findings indicated alterations in the topological properties of brain functional networks among PD-SH patients. Nevertheless, there was no statistically significant difference in the small world properties of brain networks between the PD-SH and PD-N/MH group.

As shown in the Figure 10 and Figure 11, the PD-N/MH, PD-SH, and HC groups all exhibited a sigma (σ) value greater (FDR correction, P<0.05) than 1 and a lambda (λ) value close to 1, it can be inferred that both the PD and HC groups possess small-world properties in their brain networks [53]. Compared to the HC group, the PD-SH group exhibited significantly reduced values (FDR correction, P<0.05) of C_p_ and E_loc_, while the lambda value was significantly elevated (Figure 11).

**Figure 10.**
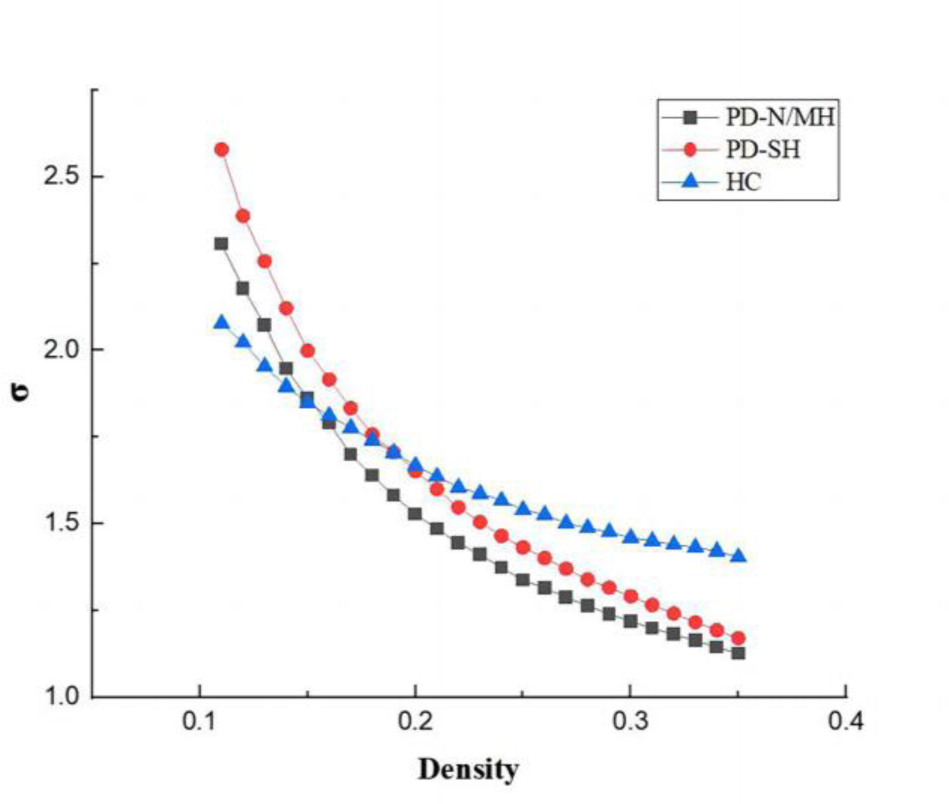
Comparison of “Small World” property among three groups.

**Figure 11.**
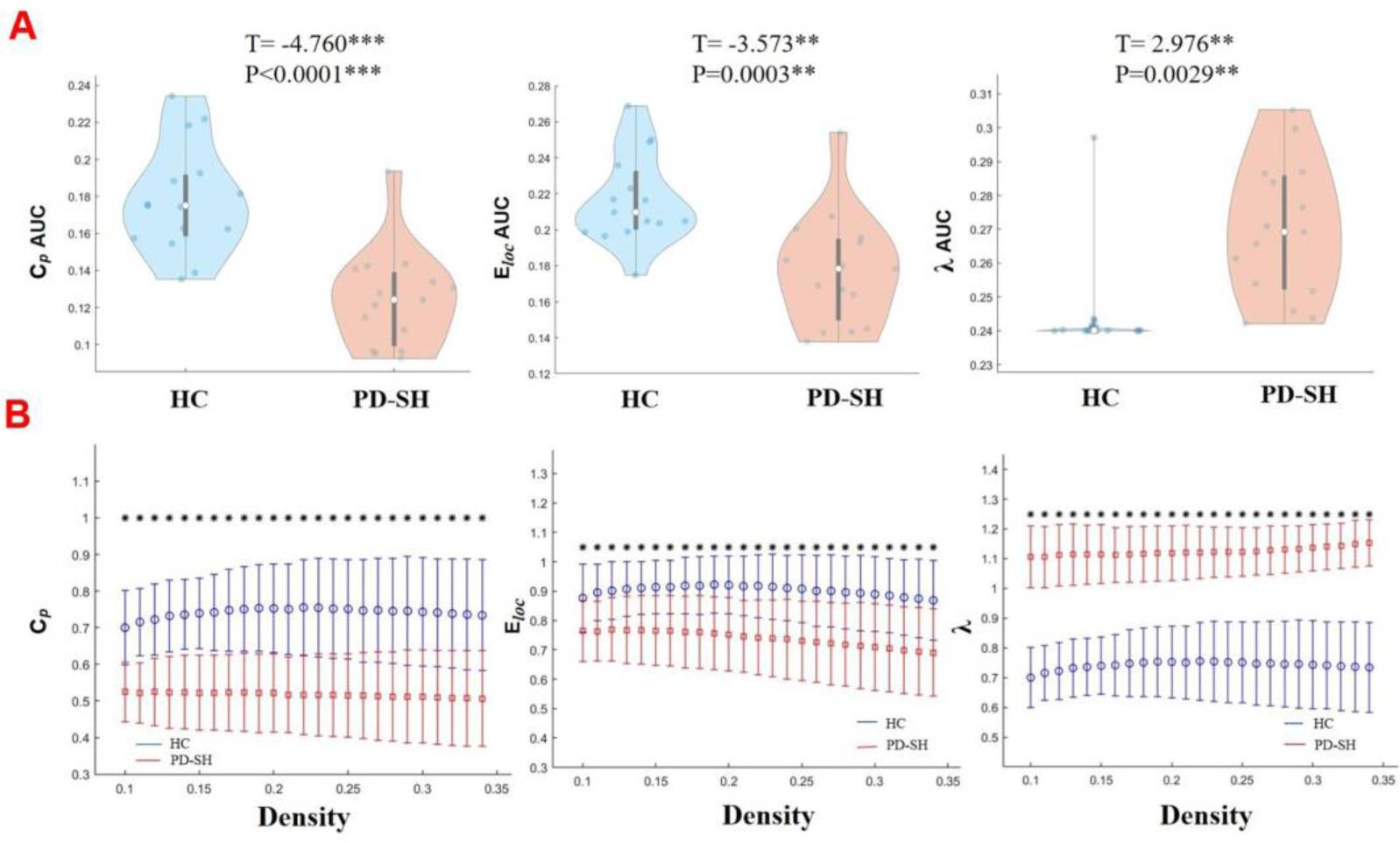
Differences in network topological properties among the PD-SH, PD-N/MH, and HC groups. **A**: Violin plots depict the distribution of mean C_p_ AUC, E_loc_ AUC and λ values, highlighting the contrast between PD-SH and HC. **B**: C_p_, E_loc_ and λvalues are shown across a density range spanning from 10% to 34%. Each point, accompanied by an error bar, represents the mean and standard deviation at specific density levels, respectively. ** denotes P <0.01; *** denotes P <0.0001.

Similarly, the PD-N/MH group also demonstrated significantly lower values of C_p_ and E_loc_ compared to the HC group, along with higher λ values (Figure 12).

**Figure 12.**
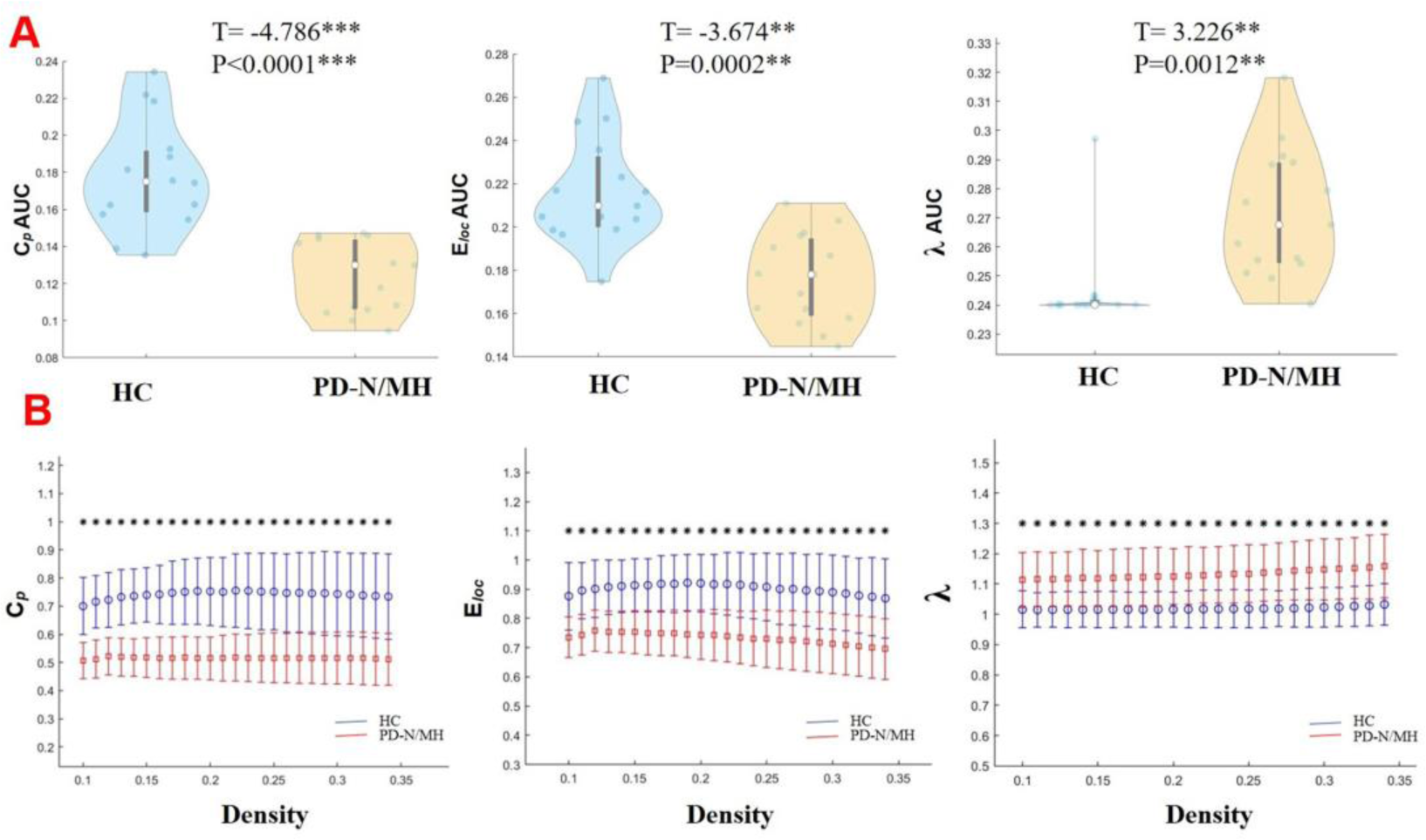
Differences in network topological properties between the HC and PD-N/MHgroups. **A**: Violin plots depict the distribution of mean C_p_ AUC, E_loc_ AUC and λ values, highlighting the contrast between PD-N/MH and HC. **B**: C_p_, E_loc_ and λ values are shown across a density range spanning from 10% to 34%. Each point, accompanied by an error bar, represents the mean and standard deviation at specific density levels, respectively. ** denotes P <0.01; *** denotes P <0.0001.

#### 3.4.2. Nodal Properties

Significant disparities were observed in the nodal characteristics between the PD group and the HC group, as detailed below: In comparison to the HC group, patients with PD-SH demonstrated a notable elevation (FDR correction, P<0.05) in the nodal betweenness centrality of the right Superior frontal gyrus, dorsolateral and right vermis; the betweenness centrality of the node located in the right Superior frontal gyrus, dorsolateral was notably elevated (FDR correction, P<0.05) in the PD-N/MH group compared to the HC group (Figure 13).

**Figure 13.**
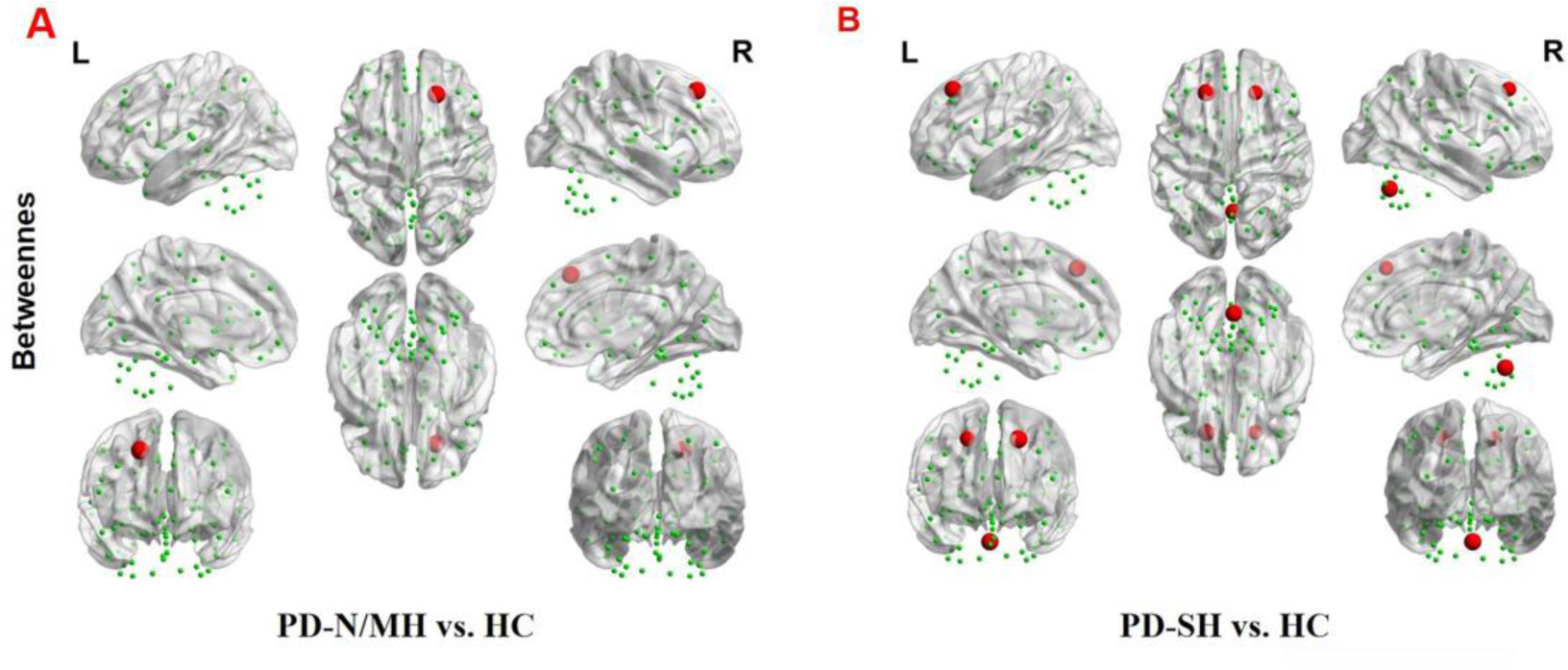
The brain regions with statistically significant difference in Betweenness centrality among PD-SH, PD-N.MH an HC group. **A**: The difference in Betweenness centrality between the PD-N/MH and HC group. **B**: The difference in Betweenness centrality between the PD-SH and HC group. The red spheres represent the brain nodes with increased betweenness centrality in PD-N/MH or PD-SH compared to HC group. R: right; L: left.

Compared to the PD-SH group, the PD-N/MH group demonstrated a notable increase (P<0.05) in betweenness centrality within the left superior cerebellum (Figure 14).

**Figure 14.**
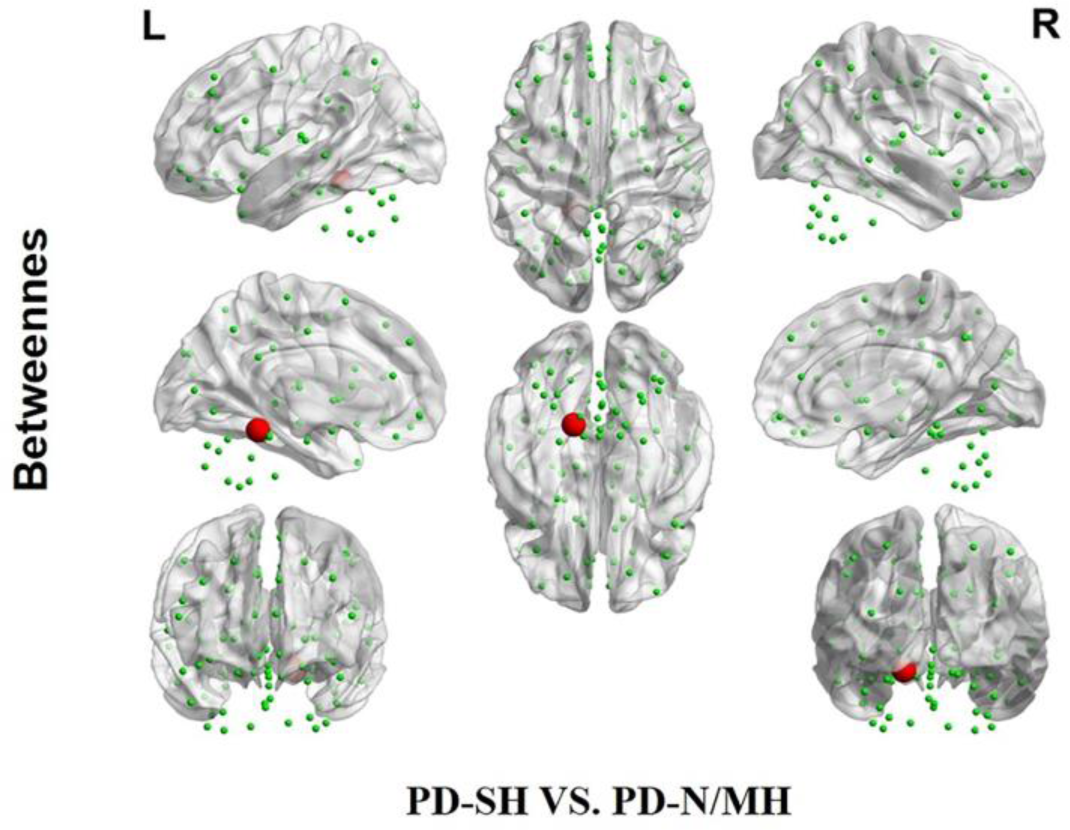
The difference in Betweenness centrality between the PD-SH and PD-N/MH group. The red spheres represent the brain nodes with increased betweenness centrality in PD-N/MH compared to PD-SH group. R: right; L: left.

We further observed a significant decrease (FDR correction, P<0.05) in nodal degree in multiple brain regions of PD-SH patients compared to the HC group, specifically the Left Inferior frontal gyrus, opercular part, left Insula, bilateral Cuneus, left Middle occipital gyrus, bilateral Inferior occipital gyrus, right Inferior parietal, but supramarginal and angular gyri, bilateral Supramarginal gyrus, bilateral Angular gyrus, bilateral Caudate nucleus, left Heschl gyrus and left Superior Cerebellum (Figure 15).

**Figure 15.**
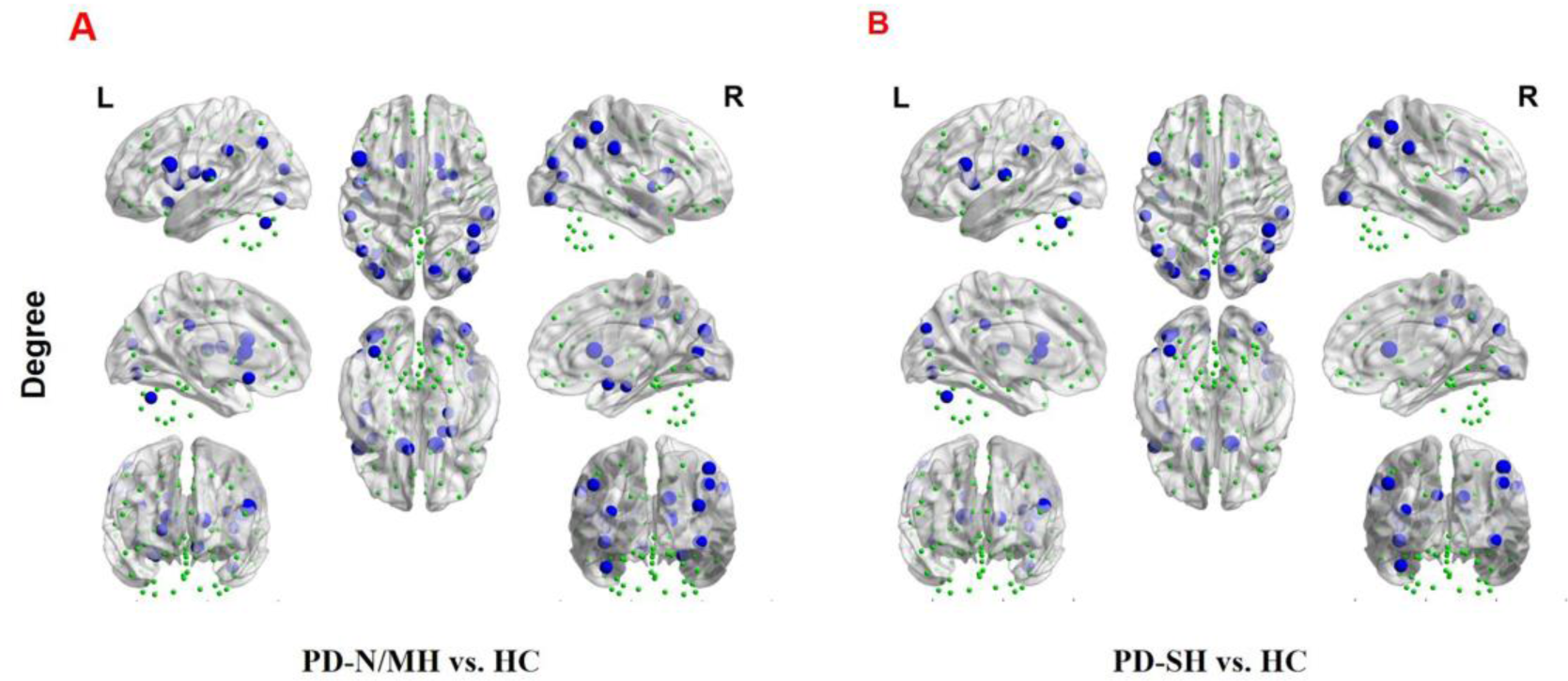
The brain regions with statistically significant difference in nodal degree among PD-SH, PD-N/MH an HC group. **A**: The difference in nodal degree between the PD-N/MH and HC group. **B**: The difference in nodal degree between the PD-SH and HC group. The blue spheres represent the brain nodes with increased nodal degree in PD-N/MH or PD-SH compared to HC group. R: right; L: left.

Compared to the HC group, the PD-N/MH group also exhibited regions with significantly reduced (FDR correction, P<0.05) nodal degree, as follows: Left Inferior frontal gyrus, opercular part, left Rolandic operculum, left Olfactory cortex, left Insula, right Parahippocampal gyrus, right Amygdala, right Calcarine fissure and surrounding cortex, right Cuneus, bilateral Middle occipital gyrus, bilateral Inferior occipital gyrus, right Inferior parietal, but supramarginal and angular gyri, bilateral Supramarginal gyrus, bilateral Angular gyrus, bilateral Caudate nucleus, rifht Lenticular nucleus, pallidum and left Superior Cerebellum.

We found that compared with PD-SH patients, PD-N/MH patients exhibited a significant decrease (P<0.05) in nodal degree in the left superior temporal lobe (Figure 16).

**Figure 16.**
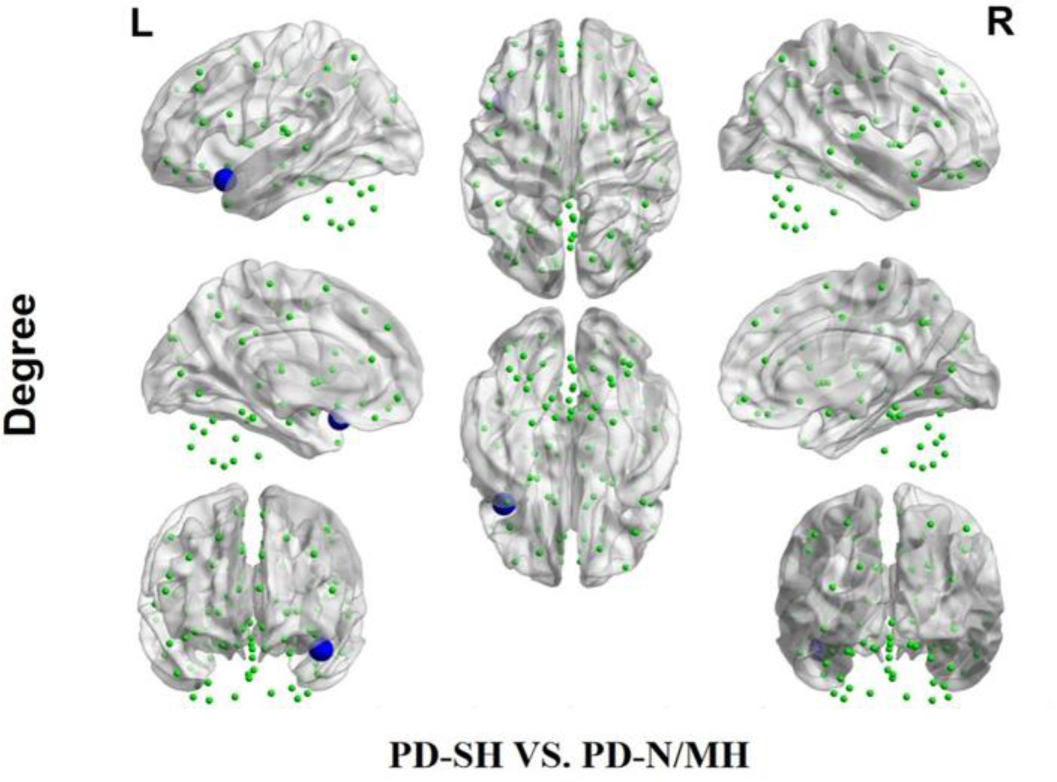
The difference in nodal degree between the PD-SH and PD-N/MH group. The blue spheres represent the brain nodes with increased nodal degree in PD-N/MH compared to PD-SH group. R: right; L: left.

### 3.5. Results of Correlation Analysis

We performed a correlation analysis to investigate the relationship between the ALFF values of patients in the PD group (PD-SH group and PD-N/MH group) and their olfactory scores (OSIT-J score), and the findings are presented below (Figure 17): the ALFF value observed in bilateral superior cerebellum, bilateral fusiform gyrus, right lingual gyrus, bilateral middle temporal gyrus, left inferior frontal gyrus, triangular part and bilateral supramarginal gyrus demonstrated a statistically significant positive correlation with the olfactory scores (R>0.5). Conversely, the ALFF values in the bilateral middle temporal gyrus, left lingual gyrus, left superior temporal gyrus, left temporal pole: superior temporal gyrus, right inferior temporal gyrus, right postcentral gyrus, left middle frontal gyrus displayed a negative correlation with olfactory scores.

**Figure 17.**
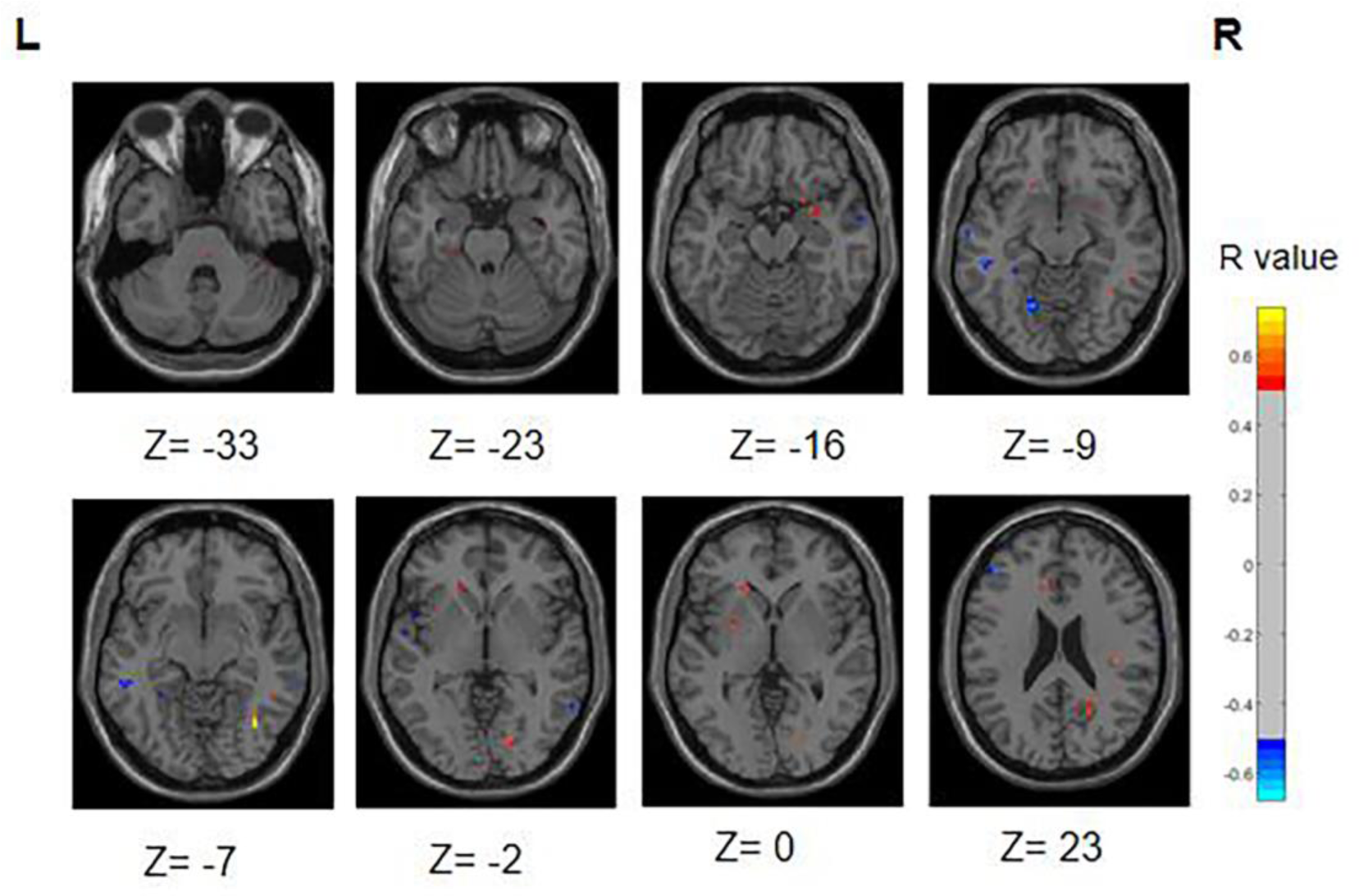
Correlation maps illustrating the relationship between ALFF values and olfactory score in PD patients. Warm colors, such as red, denote a positive correlation, while cool colors, like blue, signify a negative correlation. The color bar denotes the R-value. R: right; L: left

Similarly, we performed a correlation analysis to assess the relationship between the ReHo values and olfactory scores among patients with PD (Figure 18). The following regions demonstrated a positive correlation with olfactory scores: right inferior cerebellum, left inferior temporal gyrus, left fusiform gyrus, right inferior occipital gyrus, right superior temporal gyrus and right middle temporal gyrus (R>0.5). Conversely, the following regions right middle frontal gyrus, right superior frontal gyrus, right superior frontal gyrus, medial and right middle occipital gyrus displayed a negative correlation with olfactory scores

**Figure 18.**
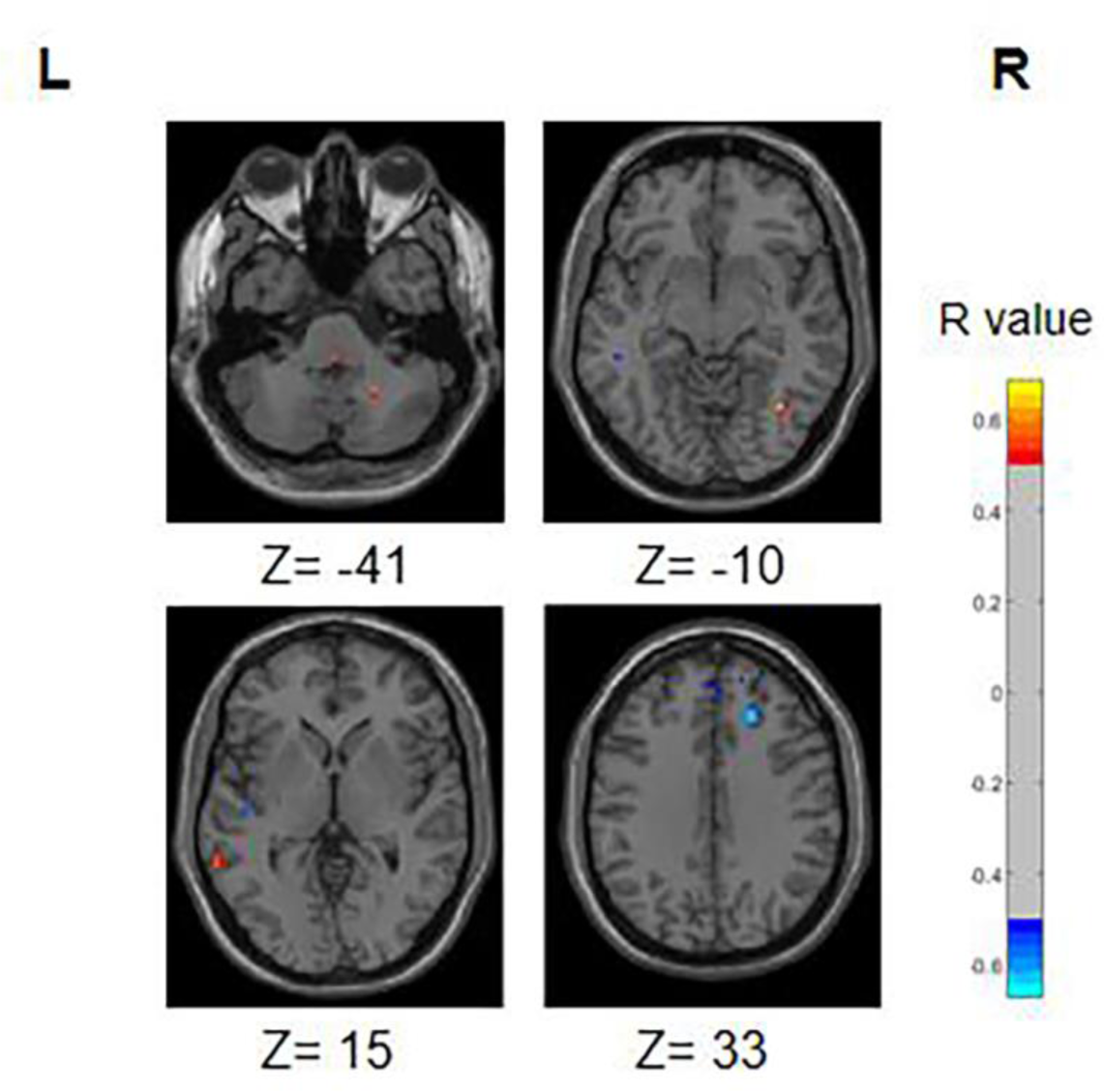
Correlation maps illustrating the relationship between ReHo values and olfactory score in PD patients. Warm colors, such as red, denote a positive correlation, while cool colors, like blue, signify a negative correlation. The color bar denotes the R-value. R: right; L: left.

During the correlation analysis between FC values and OSIT-J scores, a negative correlation (P<0.005) was observed between olfactory scores and the functional connectivity between the left superior cerebellum and right insula (Figure 19).

**Figure 19.**
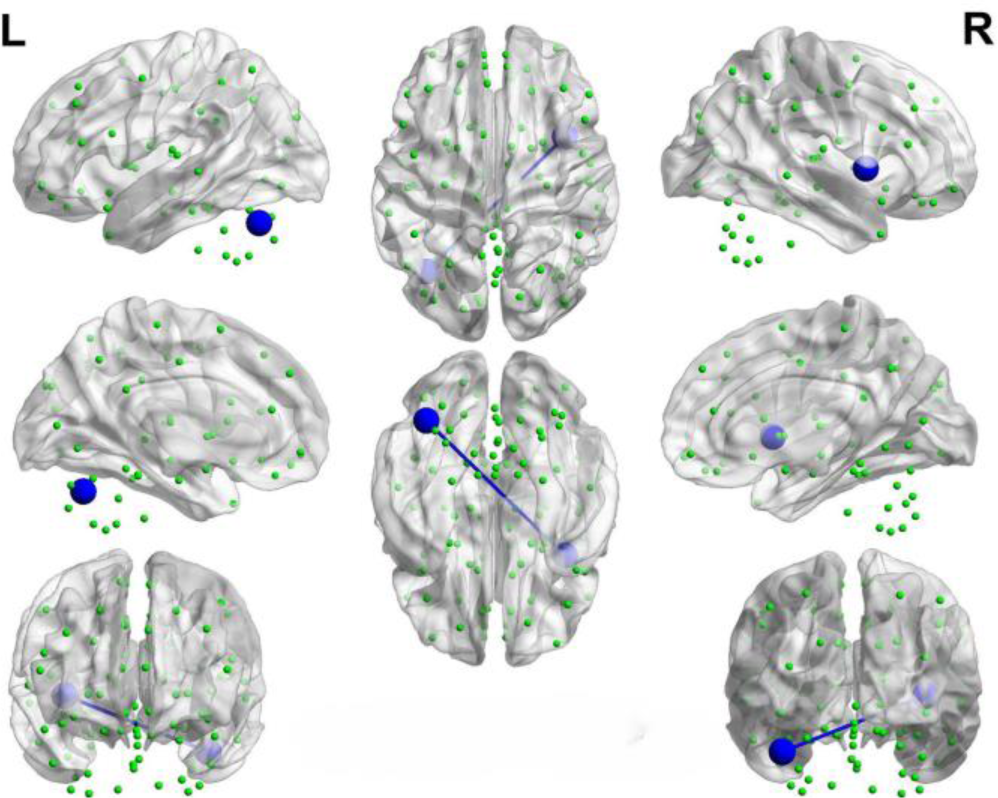
Brain map depicting functional connectivity with negative correlation. The azure sphere serves as a symbol, representing a node that corresponds to a distinct brain region, whereas the blue line denotes the existence of a negative correlation between functional connectivity and olfactory score. R: right; L: left.

We identified positive correlations between the nodal betweenness and olfactory scores in several brain regions (Figure 20), particularly the bilateral olfactory cortex, right superior occipital gyrus and left superior cerebellum (P<0.05). In addition, the nodal betweeness of right posterior cingulate gyrus was found to be negatively correlated with olfactory scores.

**Figure 20.**
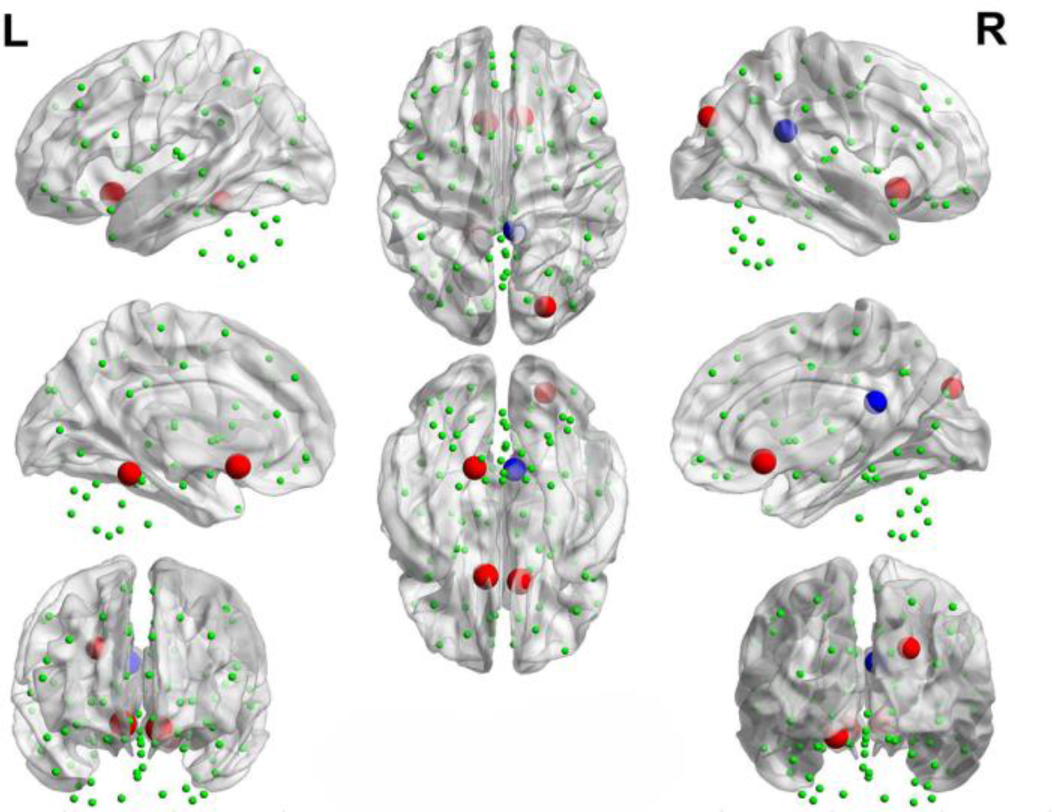
Brain map portraying nodes exhibiting correlations with OSIT-J scores. The red sphere signifies a positive correlation between the nodal betweenness and the olfactory score, whereas the blue sphere denotes a negative correlation.

However, upon analysis, it was determined that there was no significant correlation between node degree, small-world property values, and olfactory scores.

## 4. Discussion

### 4.1. Relationship between Functional Separation and Hyposmia

Functional separation serves as a tool to assess the local functional properties of brain regions, encompassing two key indicators: ALFF and ReHo [54, 55]. ALFF serves as a measure of spontaneous neural activity within local brain regions. Elevated ALFF values indicate an increase in spontaneous neural activity, whereas lower ALFF values suggest a decrease in such activity [56]. ReHo, an indicator of the synchronicity of neural activity within local brain regions [57]. In this study, we observed the existence of differential brain regions in both the PD-SH and PD-N/MH groups when compared to the ALFF values in the HC group. Notably, the PD-SH group exhibited a greater number of differential brain regions compared to the PD-N/MH group. We observed a significant elevation in ALFF values within the superior temporal gyrus in both the PD-SH and PD-N/MH groups, compared to the HC group. Furthermore, the ALFF values in the superior temporal gyrus were notably elevated in the PD-SH group in comparison to the PD-NMH group. Given the observed negative correlation between ALFF values in the superior temporal gyrus and olfactory scores, we hypothesize that the superior temporal gyrus plays a role in the pathophysiology of olfactory dysfunction in patients with PD.

In patients with Parkinson’s disease (PD), the ReHo values in the inferior cerebellum are elevated compared to those in the healthy control (HC) group. Additionally, a positive correlation exists between the ReHo values in the inferior cerebellum and olfactory scores, hinting at a potential decrease in neuronal activity within this region during olfactory dysfunction. However, the magnitude of this decrease may be modest, as no significant differences in ReHo values were detected between the PD-SH group and the PD-N/MH group in the inferior cerebellum.

The ReHo values observed in the middle temporal gyrus and superior frontal gyrus of patients with Parkinson’s disease (PD) are notably elevated compared to those in the healthy control (HC) group. Furthermore, the PD-SH group, who exhibit severe olfactory dysfunction, demonstrate significantly higher ReHo values in the middle temporal gyrus and superior frontal gyrus than the PD-N/MH group. This indicates a compensatory enhancement of neuronal activity in these regions among PD patients with severe olfactory impairment. This finding is corroborated by the negative correlation between ReHo values in the superior frontal gyrus and olfactory scores.

Significant differences in ReHo values are observed in the occipital middle gyrus between the PD-SH and PD-N/MH groups, with a negative correlation evident between ReHo values in this region and olfactory scores. This finding suggests that compensatory enhancement of local neuronal activity in the occipital middle gyrus may contribute to olfactory dysfunction.

Research has demonstrated notable gray matter atrophy in the frontal, temporal, and occipital regions among patients with Parkinson’s disease (PD) [58]. Nevertheless, our study revealed that PD patients experiencing severe olfactory dysfunction exhibited heightened brain functional activity in certain areas within these regions. This observation hints at a potential role of the frontal, temporal, and occipital regions in olfactory dysfunction among PD patients. However, further investigation is warranted to elucidate the specific molecular mechanisms involved and their interplay with neurotransmitters, including dopamine, serotonin, and acetylcholine.

### 4.2. Relationship between Functional Connectivity and Hyposmia

We observed that, in comparison to the HC group, the PD-SH and PD-N/MH groups exhibited weaker connections in the left middle frontal gyrus, the left precental gyrus, as well as between the left insula and the left caudate nucleus. Furthermore, the PD patients in both groups exhibited stronger functional connectivity compared to the HC group in the following regions: the dorsolateral and orbital parts of the left superior frontal gyrus, the connection between the medial and orbital parts of the left superior frontal gyrus, and the connection between the left inferior cerebellum and the orbital part of the left superior frontal gyrus. Based on previous research on Parkinson’s disease (PD), the accumulation of Lewy bodies in the prefrontal cortex and the disruption of the basal ganglia motor circuit represent the characteristic pathological changes associated with the condition [59]. Therefore, our findings indicate that the aberrant functional connectivity observed in the aforementioned regions is likely associated with PD itself, rather than being a consequence of olfactory dysfunction related alterations. It is worth noting that compared to the PD-N/MH group, the PD-SH group displayed significant abnormalities in functional connectivity within the cerebellum. Interestingly, the functional connectivity between the superior cerebellar lobule and the insula showed a negative correlation with olfactory scores, indicating that compensatory enhancement of cerebellar functional connectivity occurs in PD patients with olfactory dysfunction. This finding aligns with our prior research, revealing that the cerebellum and insula are linked to olfactory function in PD patients and exhibit aberrant functional connectivity with white matter fiber bundles [60]. Consequently, our study reinforces the potential significance of the cerebellum and insula in olfactory impairments among PD patients.

### 4.3. Alterations in Network Topology

In this study, using graph-theoretical analysis methods, we explored the alterations in the topological properties of brain networks in patients with PD who also had severe olfactory dysfunction. Some previous researches on the topological attributes of brain networks in PD patients have consistently shown the maintenance of small-world properties, which agrees with our own findings [61–63]. Our investigation has shown that both PD-SH and PD-N/MH possess small-world characteristics within their functional brain networks, nonetheless, notable changes have occurred in the topological attributes of these networks. When comparing PD patients with HC, it is observed that PD patients exhibit a lower clustering coefficient and local efficiency, coupled with a higher normalized characteristic shortest path length. These findings indicate that the topological structure of the brain networks in PD patients was disrupted, resulting in decreased efficiency in local information processing and integration, and a shift towards a more randomized functional brain network [64]. Certain studies have demonstrated a strong correlation between olfactory dysfunction and both local and global efficiency [65]. Perhaps the olfactory dysfunction arises from the decreased local efficiency of the brain network, which impairs the transmission of olfactory information. Despite our efforts, we did not detect any significant differences in the small-world properties of the brain between the PD-SH and PD-N/MH groups. Based on this observation, we hypothesize that the impact of olfactory dysfunction on the brain’s small-world attributes in PD patients is rather minimal, to the extent that it cannot be reliably detected in smaller sample sizes.

During the analysis of node attributes, we found significant differences in node betweenness centrality between the PD-SH and PD-NMH groups compared to the HC group. Notably, the PD-SH group exhibited a larger number of nodes with such differences. Additionally, the PD-NMH group showed a significant increase in nodal betweenness centrality specifically in the superior cerebellar lobule compared to the PD-SH group. Intriguingly, a positive correlation was observed between the betweenness centrality of the superior cerebellar lobule and olfactory scores. These observations strongly suggest that a decrease in betweenness centrality in the superior cerebellar lobule may contribute to the pathogenesis of olfactory dysfunction.

Significant differences in node degree within the superior temporal gyrus have been observed between the PD-SH and PD-N/MH groups. Despite the absence of a significant correlation between these node degree differences and olfactory scores, we cannot conclusively dismiss the potential role of the superior temporal gyrus in olfactory dysfunction. This lack of correlation may be attributed to statistical errors arising from a limited sample size.

### 4.4. Limitations

Firstly, the data utilized in this study was sourced from public databases. However, in the future, it is imperative that we personally gather data to enhance its precision and reliability. Secondly, this study adopts a cross-sectional and retrospective approach, which poses a challenge in determining whether olfactory dysfunction precedes changes in brain function imaging or whether modifications in brain function precede the observed imaging manifestations. Consequently, it is imperative to conduct longitudinal and prospective studies in the future, encompassing follow-up research on PD-N/MH patients, to further clarify these relationships. Finally, it is noteworthy that task-based functional magnetic resonance imaging (fMRI) exhibits enhanced sensitivity and accuracy in detecting brain function changes. Consequently, in the future, it is advisable to consider utilizing fMRI examinations that involve olfactory stimulation to delve deeper into the mechanisms responsible for olfactory dysfunction in PD patients.

## 5. Conclusion

Utilizing indices such as ALFF, REHO, FC, and brain network topological properties, this study revealed significant differences in brain function among patients categorized into the PD-SH, PD-N/MH, and HC groups, specifically with regards to their brains’ functional segregation and integration capabilities.

From the lens of local brain function, we speculate that alterations in the functional activities within the temporal, frontal, occipital lobes, and the cerebellum contribute significantly to the olfactory impairment observed in patients with Parkinson’s disease. From the angle of brain functional connectivity, the presence of abnormal connectivity between the cerebellum and the insula, as well as internally within the cerebellum, could potentially contribute to the olfactory impairment observed in patients with Parkinson’s disease. Additionally, in terms of brain network topological properties, despite sharing small-world properties, notable differences in node degree within the superior temporal gyrus and the betweenness centrality of the superior cerebellar lobule between the PD-SH and PD-N/MH groups indicate potential associations between alterations in nodal characteristics of these regions and olfactory dysfunction. Based on these comprehensive analyses, we hypothesize that these regions play a crucial role in the pathogenesis of olfactory dysfunction observed in PD patients.

## AUTHOR CONTRIBUTIONS

LG and WC equally contributed to the study. LG and WC contributed to the draft, and analyzed the data. LG, WC, NW and YS conceived this study together. JZ, HY, and JW contributed to the collection of clinical and fMRI data. YS and NW strictly revised the manuscript. All authors have read and agreed to the published version of the manuscript.

## ACKNOWLEDGMENTS

We would like to thank all the patients for their participation in this project.

## DATA AVAILABILITY STATEMENT

Publicly available datasets were analyzed in this study. This data can be found here: The data was obtained from the OpenfMRI database. Its login number is ds000245 (https://www.openfmri.org/dataset/ds000245/).

